# Spatial modeling of extinct Southeast Asian Hobbit *Homo floresiensis* habitat suitability based on its diet availability and climatic parameters as a proxy

**DOI:** 10.1101/2021.09.06.459154

**Authors:** Andri Wibowo

## Abstract

The discovery of a new prehistoric species of the genus *Homo* remains a significant matter of intense interest. One of significant discovery recently is the *Homo floresiensis,* representing a small-bodied and small-brained hominin, excavated, and found in Liang Bua Cave in Flores Island, East Nusa Tenggara, Indonesia. This species height was only about 106 cm (3’6“) and the weight was 30-40 kg (66-86 lbs). *H. floresiensis* was known consumed extant murine rodents as its diets as this was evidence found in Liang Bua Cave. Then this study aims to model the *H. floresiensis* suitable habitat using maximum entropy method and 2 extant murine rodents, *Rattus exulans* and *R. rattus* as a proxy. The results show that the most suitable habitats for *H. floresiensis* indicated by suitable habitat values close to 1 were concentrated in the central of Flores Island that was overlapped with mountainous areas with elevation ranging from 1500 to 2000 m. These suitable habitats were also overlapped with dense vegetation covers, volcanic rock, and Kiro rock formation. Climatic parameters that limit the distributions of *H. floresiensis* were annual mean temperature, isothermality, minimum temperature of coldest period, and precipitation seasonality. Parts of Flores Island with the low temperature below 20 °C were favorable for *H. floresiensis* while an increase in isothermality limits the *H. floresiensis* distributions.

## Introduction

Hobbit is a nickname of hominin species, identified as *Homo floresiensis* excavated and found from Flores Island or administratively was under East Nusa Tenggara Province, Indonesia. This species was formerly known as a microcephalic dwarf modern human being (Colin 2007, Baab 2012) and assumed as an ancestor of Rampasa tribe that still living today in Flores Island. Whereas recent studies confirmed that *H. floresiensis* most closely resembles hominin species from the Late Pliocene and Early Pleistocene including species of *H. habilis*, *H.ergaster*, and *H. georgicus* (Argue et al. 2006).

In Indonesia, Flores is one of many Wallacean islands located on east of Wallace’s Line and west of Lydekker’s Line. Wallacean islands are important for biogeography sciences since it has been connected via land bridges to either the Asian continent to the west or the Greater Australian continent to the east. This longstanding separation from the surrounding continents either with Australia or Asia has severely limited the ability of species mainly animals to disperse either into or away from the Wallacean islands. As a result, Flores Island was isolated and in Flores there were only a small number of mammal and reptile species during the entire Pleistocene. Those species included komodo dragons and other smaller monitor lizards, crocodiles, several species of Stegodon, (an extinct close relative of modern elephants), giant tortoise, and several kinds of small, medium, large-bodied rats, and *H. floresiensis* (Bergh et al. 2001).

Recently, the diets of *H. floresiensis* have been identified. Fossils of animals were excavated and found nearby *H. floresiensis* fossils were found. Excavation in Liang Bua resulted in 230,000 bone fragments (Hafsari 2017) that were identified belongs to murine rodents under Rodentia Order, Muridae Family, and Murinae Subfamily or known as rats (Veatch 2014). At least there were 6 rat Generas including *Papagomys* with species of *P. armandvilley* and *P. theodorverhoeveni*, *Spelaeomys florensis*, *Hooijeromys nusatenggara*, *Komodomys rintjanus*, *Paulamys naso*, and *Rattus* including *Rattus hainaldi* and *Rattus exulans*. Humerus and femur of medium (100-300 g) followed by small (<100 g) murine rodents (Veatch et al. 2019) were accounted for almost half of the fossils found in Liang Bua. Small murine was identified as *Rattus hainaldi* and *Rattus exulans* and medium sized was *Rattus rattus*. Hafsari (2017) argued that the presence of murine rodents in Liang Bua due to consumption of murines by cave occupants in this case was presumably *H. floresiensis*. Even at the current time, native people nearby Liang Bua still consume rats. Despite *H. floresiensis* has been extensively studied, information on where did these hominin species was previously distributed was still limited. Here this study aims to estimate the suitable habitat of extant *H. floresiensis* using its diet, a murine rodent species, distribution, as a proxy using predator prey interaction distribution model (Trainor et al 2014).

## Methodology

### Study area

The location was a Flores Island (Figure 1) located in the eastern part of Indonesia. This island has a size of 13,540 km^2^ and has geographical coordinate of 120° and 122° longitude and 8.3° and 8.7° latitude. The highest point was 2,370 m located in the west part of the island and observed in mount Poco mandasawu. The landscape of Flores Island was a combination of low land and mountainous region with hilly landscape. The climate of Flores Island is tropical dry with a fairly long dry season, which is about 8 months per year with uneven distribution of rainfall. The air temperature varies between 21.2 °Celsius - 33.4 °Celsius.

**Figure 1.**
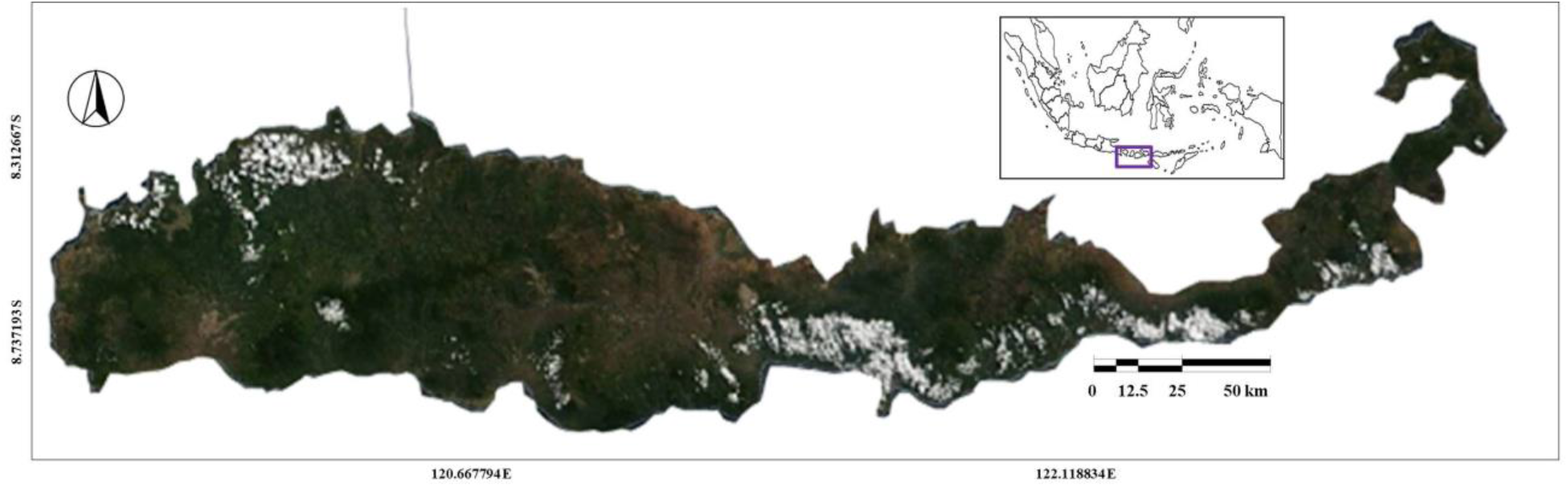
Study area in Flores Island, East Nusa Tenggara, Indonesia

### Murine data retrieval

This study used the presence data of 2 murine rodent species identified as the potential *H. floresiensis* diet. Those murine species included extant *Rattus exulans* and *Rattus rattus*. The presence and locality data of those species were retrieved from Global Biodiversity Information Facility (Boitani et al. 2011, Prieto-Torres & Pinilla-Buitrago 2017, Truong et al. 2017). The retrieved data included the geographical coordinate of those murines recorded in Flores island. The coordinates including longitude and latitude then mapped using GIS to be used for further distribution modeling analysis.

### Elevation, geology, and tree cover data retrieval

This study uses several environmental data including elevation, geology, and tree cover thematic data. The elevation data of Flores Island denoted in m was retrieved from Shuttle Radar Topography Mission (Patel & Sarkar 2010). The geological data including rock formation and aquifer lithology were obtained from the Geological Agency of Indonesia. The tree covers data indicating the presence of forest in Flores Island were retrieved using multispectral satellite imagery from the Landsat 7 thematic mapper plus (ETM+) sensor (Hansen et al. 2013).

### Murine distribution modeling

The murine distribution was estimated using murine presence data retrieved from GBIF and mapped using GIS as described in previous steps. Distribution estimation of species was developed followed methods by Zhang et al. (2019). The potential murine distribution was modeled using the principle of maximum entropy to calculate the most likely distribution of the murine species in the function of occurrence localities and environmental variables. Suitability habitat was classified and ranked from 0 as the least suitable to the 1 for the most suitable habitat. The model used 15 climatic variables (Beaumont et al. 2005) retrieved from WorldClim database. The complexity of the model was tested using the Akaike information criterion values (AICc) corrected under different parameter conditions. AIC is a standard to measure the goodness of statistical model fitting and it generally gives priority to the parameters with small AIC values for simulation

## Results and Discussions

*Rattus exulans* and *Rattus rattus* were 2 murine species that were still living across Flores Island landscape. Based on the GBIF records, there were 5 locations where both murine species occurrences were overlapped. While *R. rattus* presence distribution was wider since this species was also found in the most western parts of the island (Figure 2.a.). Both species have a wide distribution from east to central and west of Flores Island.

**Figure 2.**
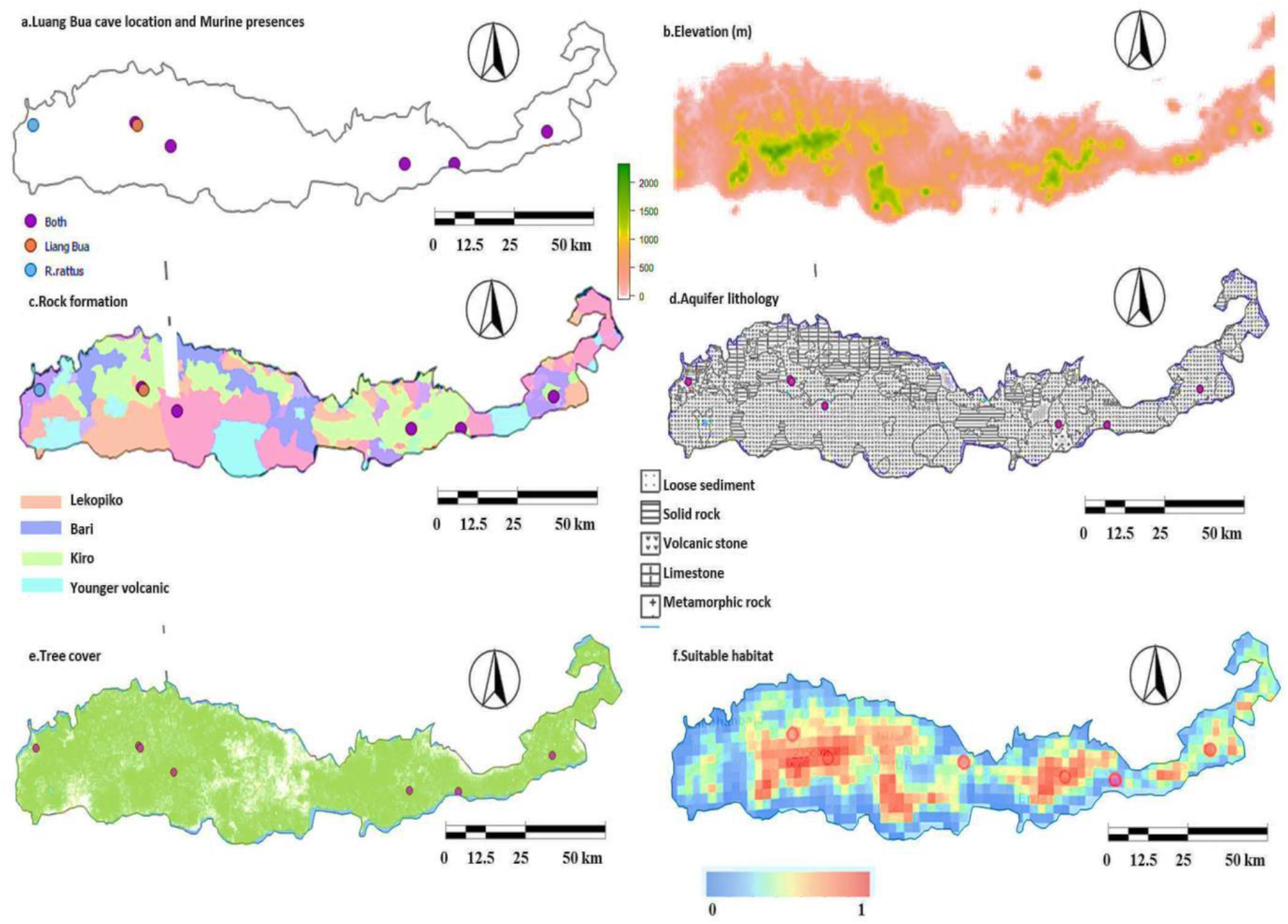
Rotate clock wise: (a) Liang Bua cave location (orange dot) and Murine presences in Flores Island; (b) Elevation (m) of Flores Island; (c) Rock formation of Flores Island; (d) Aquifer lithology of Flores Island; (e) Tree covers of Flores Island; (f) Estimated habitat suitability of Southeast Asian Hobbit based on Murine species (*R. rattus* and *R. exulans*) estimated species distribution modeling.

The landscape of Flores Island was mostly dominated by a hilly landscape that was very common in west and central parts, and some patchy hilly landscapes in the east part. *R. exulans* and *R. rattus* occurrences and Liang Bua Cave location at present time were overlapped with the hilly landscape. Both murines were occupying hilly landscapes in the west and east parts of the island (Figure 2.b.). Flores Island was comprised of several rock formations. In this island, Liang Bua was located within Kiro formation and surrounded by Lekopiko, Bari, and younger volcanic formations. Correspondingly, both murines were also presented within Kiro formation (Figure 2.c). Kiro formation on Flores Island was characterized by deposits of volcanic materials (Figure 2.d) and breccia rocks (Suharji et al. 2013) and this indicates that this rock formation was shaped and formed through a volcanic process rather than tectonic process (Hartono 2011). Volcanic land was suitable for species since volcanic materials contain substances that can increase land fertility and support vegetation that can be consumed by murines. Rats are among species that first colonize and more adapt to the volcanic land ecosystems (Thornton et al. 2001).

Most Flores Island still has dense vegetation covers (Figure 2.e). Whereas central parts of the island were the only parts that have less covers. Murine species consumed by *H. floresiensis* were preferring mostly habitats with tree covers. While *R. rattus* has wider tolerance since this species was also found in open areas with less tree covers in the most western parts of the island where *R. exulans* was not present in here.

Figure 2.f presents the suitable habitats for murine rodents that may represent the suitable habitats for *H. floresiensis* in Flores Island. The suitable habitats were estimated based on maximum entropy analysis. It was estimated that most west, central, and east parts of Flores Island mainly in central parts were estimated suitable for murines and presumably its pedators, the *H. floresiensis*. Coastal parts were the only areas that were not suitable. The highest parts and the most suitable habitats indicated by suitable habitat values close to 1 were concentrated in the central of Flores Island that was overlapped with mountainous areas with elevation ranging from 1500 to 2000 m.

The results of the maximum entropy indicated the climatic parameters (Figure 3) that contributed more to the distribution and habitat suitability of murines were annual mean temperature, isothermality, minimum temperature of coldest period, and precipitation seasonality. Murine habitat suitability was having linear correlation with those parameters. Parts of Flores Island with the low temperature below 20 °C were favorable for murines while an increase in isothermality parameter limits the murine distributions.

**Figure 3.**
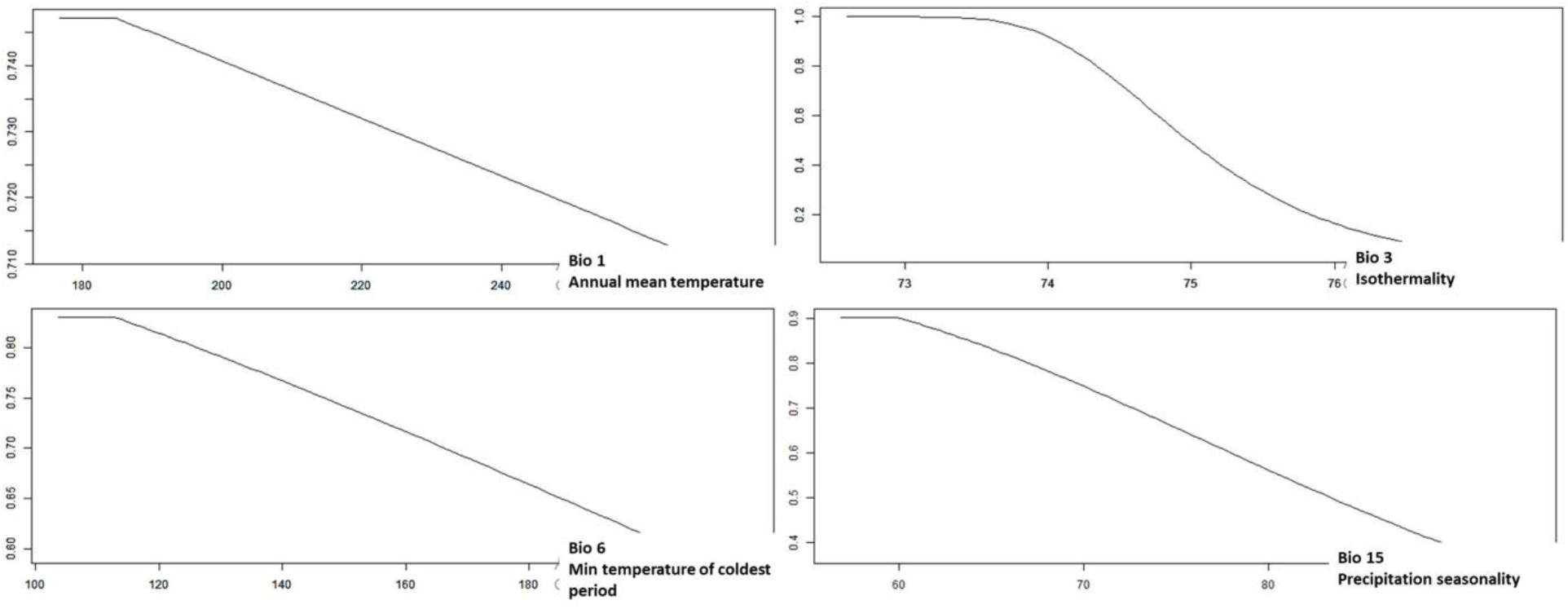
Climatic parameters affecting the habitat suitability and distributions of *R. rattus* and *R. exulans* in Flores Island

## Conclusion

Two extant murine rodents, *R. rattus* and *R. exulans* that formerly were consumed by extinct *H. floresiensis* can be used as a proxy to estimate the suitable habitats for *H. floresiensis* in Flores Island. The modeled suitable habitats were overlapped with high elevation, vegetation covers, and Kiro rock formations. High annual mean temperature, high isothermality, and high precipitation seasonality were climatic parameters that limit the distribution and suitable habitats of *R. rattus* and *R. exulans* and presumably their predators, the *H. floresiensis.*

